# Enrichment analysis on regulatory subspaces: a novel direction for the superior description of cellular responses to SARS-CoV-2

**DOI:** 10.1101/2021.12.15.472466

**Authors:** Pedro Rodrigues, Rafael S. Costa, Rui Henriques

## Abstract

**Statement:** The enrichment analysis of discriminative cell transcriptional responses to SARS-CoV-2 infection using biclustering produces a broader set of superiorly enriched GO terms and KEGG pathways against alternative state-of-the-art machine learning approaches, unraveling novel knowledge.

**Motivation and methods:** The comprehensive understanding of the impacts of the SARS-CoV-2 virus on infected cells is still incomplete. This work identifies and analyses the main cell regulatory processes affected and induced by SARS-CoV-2, using transcriptomic data from several infectable cell lines available in public databases and in vivo samples. We propose a new class of statistical models to handle three major challenges, namely the scarcity of observations, the high dimensionality of the data, and the complexity of the interactions between genes. Additionally, we analyse the function of these genes and their interactions within cells to compare them to ones affected by IAV (H1N1), RSV and HPIV3 in the target cell lines.

**Results:** Gathered results show that, although clustering and predictive algorithms aid classic functional enrichment analysis, recent pattern-based biclustering algorithms significantly improve the number and quality of the detected biological processes. Additionally, a comparative analysis of these processes is performed to identify potential pathophysiological characteristics of COVID-19. These are further compared to those identified by other authors for the same virus as well as related ones such as SARS-CoV-1. This approach is particularly relevant due to a lack of other works utilizing more complex machine learning tools within this context.

## 1. Introduction

The infection of humans by Severe Acute Respiratory Syndrome Corona Virus 2 (SARS-CoV-2) represents a major global health concern, with deaths having surpassed 5.3 million according to the World Health Organization^1^. Worldwide initiatives to publicly share data related to the virus provides an opportunity to draw novel insights into the infectious disease, which has enabled continuous breakthroughs in the understanding of how the virus can enter and utilize the cellular machinery to replicate itself and infect other cells. The knowledge relating to these mechanisms has been pushed forward mainly by a generic understanding of the process of viral replication, the transcriptomic properties of the virus, and the study of differentially expressed genes after infection and subsequent comparison to ones affected by other viral strains. This later line of research is powered by the sequencing of RNA transcripts in infectable cell lines, chosen according to the level of permissivity to infection, as well as cells collected from organisms susceptible to infection, such as humans and ferrets [1].

Despite the ongoing breakthroughs, the regulatory responses to SARS-CoV-2 infection are still not comprehensively known [2]. For instance, the role played by genes with moderate differential expression, or how interactions between multiple genes support or prevent viral replication, are still being actively updated [3]. In addition, most works in this field do not explore the role of machine learning approaches, such as clustering, predictive modeling and biclustering, to aid in the identification of differentially expressed genes and related biological processes [4].

This work aims to assess to which extent can clustering, predictive modeling and biclustering stances aid the identification of biological functions and pathways from transcriptomic data. In addition, a comprehensive analysis of the produced processes and pathways using these stances is undertaken to better understand the viral life-cycle and interactions with the cell, as well as the defence mechanisms employed by the cell against the virus.

The role of different machine learning approaches to produce gene sets for enrichment analysis is experimentally compared, showing that biclustering stances significantly aids the knowledge acquisition process. In particular, the recent class of pattern-based biclustering approaches show distinctive ability to produce a comprehensive set of superiorly enriched biological annotations in well-established knowledge bases.

The manuscript is organized as follows: section 2 covers related contributions; section 3 explores the datasets; section 4 presents the proposed methodology; section 5 experimentally compares the role of state-of-the-art machine learning stances to aid enrichment analysis, together with the description of the identified biological processes. Finally, major concluding remarks are drawn.

## 2. Related Work

Blanco-Melo et al. [1] profiled the transcriptional response of cells to infection by SARS-CoV-2 and other respiratory viruses, including RSV, IAV and HPIV3 from data collected by the authors and MERS-CoV and SARS-CoV-1 from data collected in [5]. The cells analysed consisted in three main groups: respiratory cell lines, including NHBE, A549 and Calu-3 cells; human respiratory tract cells extracted from infected and non-infected individuals; and cells extracted from infected and non-infected ferrets. The second and third groups were used to ascertain if the gene signatures matched the ones found *in vitro*. Additionally, the authors treated cells with universal IFN*β* to determine whether or not SARS-CoV-2 is sensitive to IFN-I. The treatment resulted in highly decreased viral replication, confirming the hypothesized sensitivity. To investigate how infection affects the cell transcriptome, the authors performed a differential expression analysis on NHBE cells, which revealed significant differences between the response to infection by SARS-CoV-2 and other viral strains. Functional enrichment was further performed on the differentially expressed genes to better understand the cellular functions affected by SARS-CoV-2 infection. The main factors consistent throughout the various models tested was the production of cytokines and the corresponding transcriptional response, as well as the induction of a subset of interferon stimulated genes (ISGs).

Ochsner and co-workers [6] analyzed multiple datasets to better identify the transcriptional response of human cells to SARS-CoV-2 infection as well as comparing it with MERS-CoV, SARS-CoV-1 and IAV in order to identify possible common impacts between viral strains. The authors generated consensomes by analysing how frequently the corresponding genes were differentially expressed throughout the various datasets. Similarly to Blanco-Melo et al., the authors found ISGs to have significant induction levels.

Wyler et al. [7] performed a comprehensive analysis of the transcriptional response of three cell lines, Caco-2 (a gut cell line), Calu-3 and H1299 (both lung cell lines). The authors began by identifying the susceptibility of each cell line to SARS-CoV-2 infection, which revealed H1299 cells had the lowest percentage of viral reads. Caco-2 and Calu-3 cells had comparable levels, despite the latter revealed visible signs of impaired growth and cellular death, as opposed to the former. Additionally, Calu-3 cells showed a strong induction of interferon-stimulated genes, with cytokines among these, in agreement with the findings of others.

In [8], the authors performed a genome-wide CRISPR screen on an African green monkey cell line (Vero-E6), a method used for identifying genes or genetic sequences with a target physiological effect, in this case aiding (proviral) or preventing (anti-viral) infection. To this end, surviving cells from populations infected with SARS-CoV-2 were harvested 7 days post-infection. A genome-wide screen was performed and a z-score applied to identify genes associated with increased or decreased resistance to SARS-CoV-2-induced cell death. The gene with the strongest pro-viral effect was ACE2, the protein facilitating viral entry into the cell. TMPRSS2, another gene posited to play a role in the entry of SARS-CoV-2 into the cell, was not identified significantly as pro or anti-viral, whereas the CTSL gene, which encodes the Cathepsin L protease and can also play a role in viral entry, was identified as pro-viral.

Due to thrombotic complications being common among COVID-19 patients, Manne et al. [9] investigated the functional and transcriptional changes elicited by SARS-CoV-2 infection in platelets. The data showed that SARS-CoV-2 infection does indeed alter the platelet transcriptome. To detect these changes, when comparing two groups with normal distributions, a paired t-test was used and when comparing two groups with non-normal distributions a Mann-Whitney test was used, considering a two-tailed *p*-value *<* 0.05 as statistically significant. COVID-19 further induces functional and pathological changes to platelets, including thrombocytopenia (abnormally low numbers of platelets), despite the platelets not presenting detectable levels of ACE2. This may be a contributing factor to the pathophysiology of COVID-19.

In [10], the authors tested the pathogenesis of the SARS-CoV-2 virus on transgenic mice presenting the human ACE2 gene. The infection of these mice by SARS-CoV-2 resulted in high mortality rates, especially in male mice. The transcriptional analysis of the lungs of infected animals revealed increases in transcripts involved in lung injury and inflammatory cytokines, in agreement with findings in humans.

Though there are multiple authors applying machine learning and more complex statistical models to COVID-19 patient biometric data, in order to analyse the characteristics and the outcome of the disease, these approaches have been more scarcely applied to transcriptomic data. The objective of this work is to fill this gap, addressing the question of whether the application of these approaches to this data can assist function enrichment analysis, yielding novel insights into the disease.

## 3. Dataset

In order to assess the proposed approach, we used the transcriptomic data (RNA-Seq) taken from the literatura (Gene Expression Omnibus, GEO accession GSE147507 ^2^) [1]. A schematic of the structure of the publicly available dataset is presented in Figure 1. The samples are divided into different *series* (a subset of samples), each comprising the behavior of a single cell line among different sets of experimental conditions. These also correspond to particular experiments being run, with each experiment containing multiple replicas of each experimental condition being tested. As such, the assumed independence between replicas is an important factor to test, since being able to use samples from multiple experiments simultaneously could significantly increase the amount of data available, and thus improve the reliability of the analysis.

**Figure 1:**
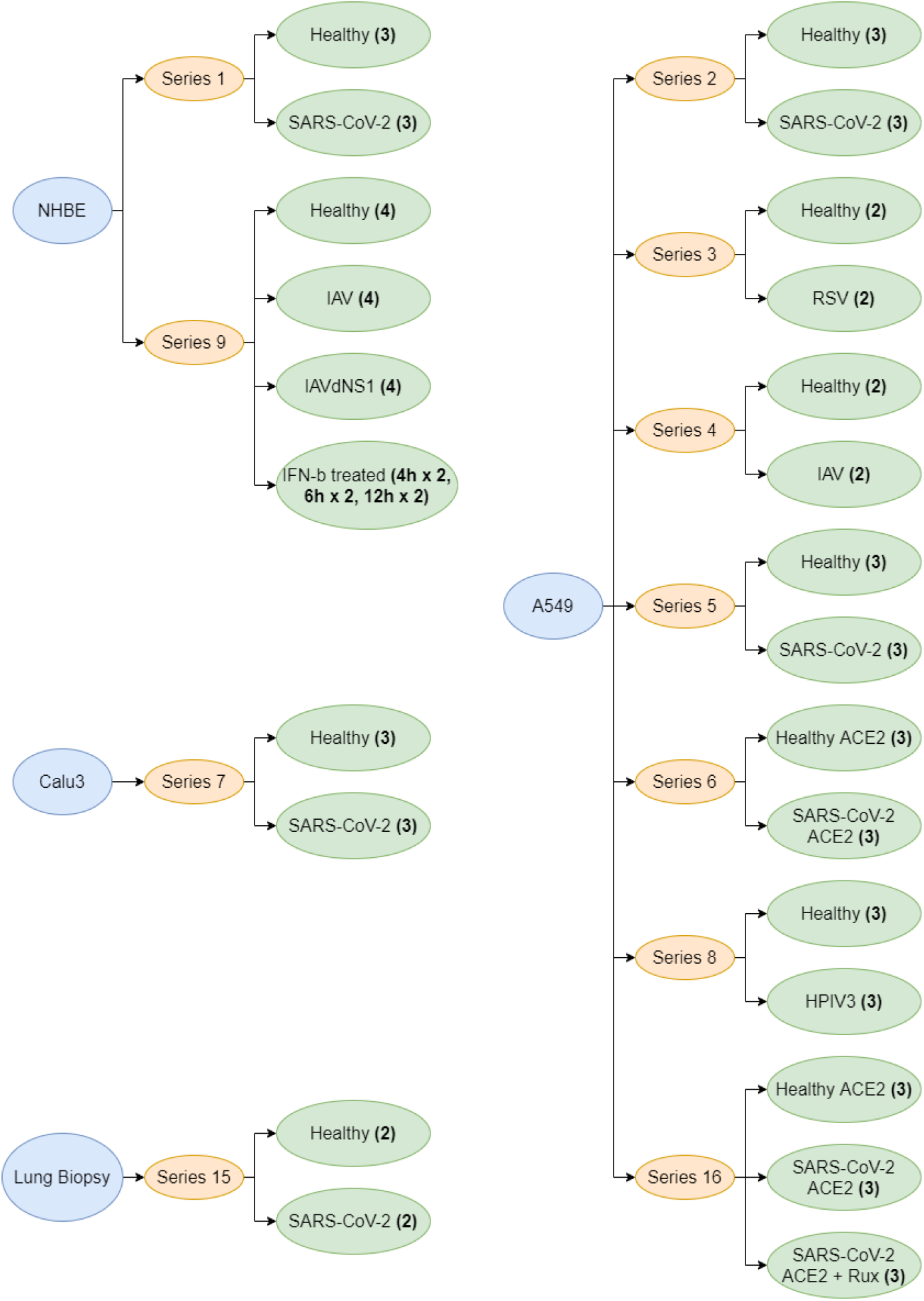
Overview of the structure of the dataset used in this study. Numbers between parentheses represent the number of data points.

For NHBE (normal human bronchial epithelial) cells, there are a total of 9 samples of healthy cells (3 belonging to *series* 1 and 4 to *series* 9), 3 samples of SARS-CoV-2 infection (all part of *series* 1), 4 samples of IAV infection (all in *series* 9), 4 samples of infection by an IAV strain which lacks the NS1 protein and, finally, 2 samples of cells treated with IFN*β* 4, 6 and 12 hours post treatment. For A549 (adenocarcinomic human alveolar basal epithelial) cells, there are 13 samples of healthy cells (3 each of *series* 2, 5 and 8, 2 each of *series* 3 and 4), 6 samples of SARS-CoV-2 infection (3 each of *series* 2 and 5), 2 samples of IAV infection (*series* 4), 2 samples of RSV infection (*series* 3) and 3 samples of HPIV3 infection (*series* 8). Blanco-Melo et al. [1] noted A549 cells had low viral counts, which was posited, in agreement with others, to be due to the low expression of ACE2 in these cells. Thus, data of A549 cells with added ACE2 (A549-ACE2) was also made available. In particular, 6 samples of healthy cells (3 each of *series* 6 and 16), 6 samples of cells infected by SARS-CoV-2 (3 each of *series* 6 and 16) and, finally, 3 samples of cells after treatment with Ruxolitinib (*series* 16). For Calu3 cells (generated from a bronchial adenocarcinoma), there are 3 samples of healthy cells and 3 samples of cells infected by SARS-CoV-2 (all belonging to *series* 7). There are an additional 2 samples from a lung biopsy of two healthy human donors (one male, one female), as well as 2 samples from a single deceased male patient of COVID-19.

Since the original data is highly skewed, the datasets were adjusted by a log2-transform for all subsequent analysis. Figure 2 depicts the distribution of expression levels among *in-vitro* cell lines and lung biopsy was then undertaken. The standard deviation of gene expression within healthy and infected cells was subsquently computed to preliminarily verify if there are significant differences between healthy and infected cells (Figure 3).

**Figure 2:**
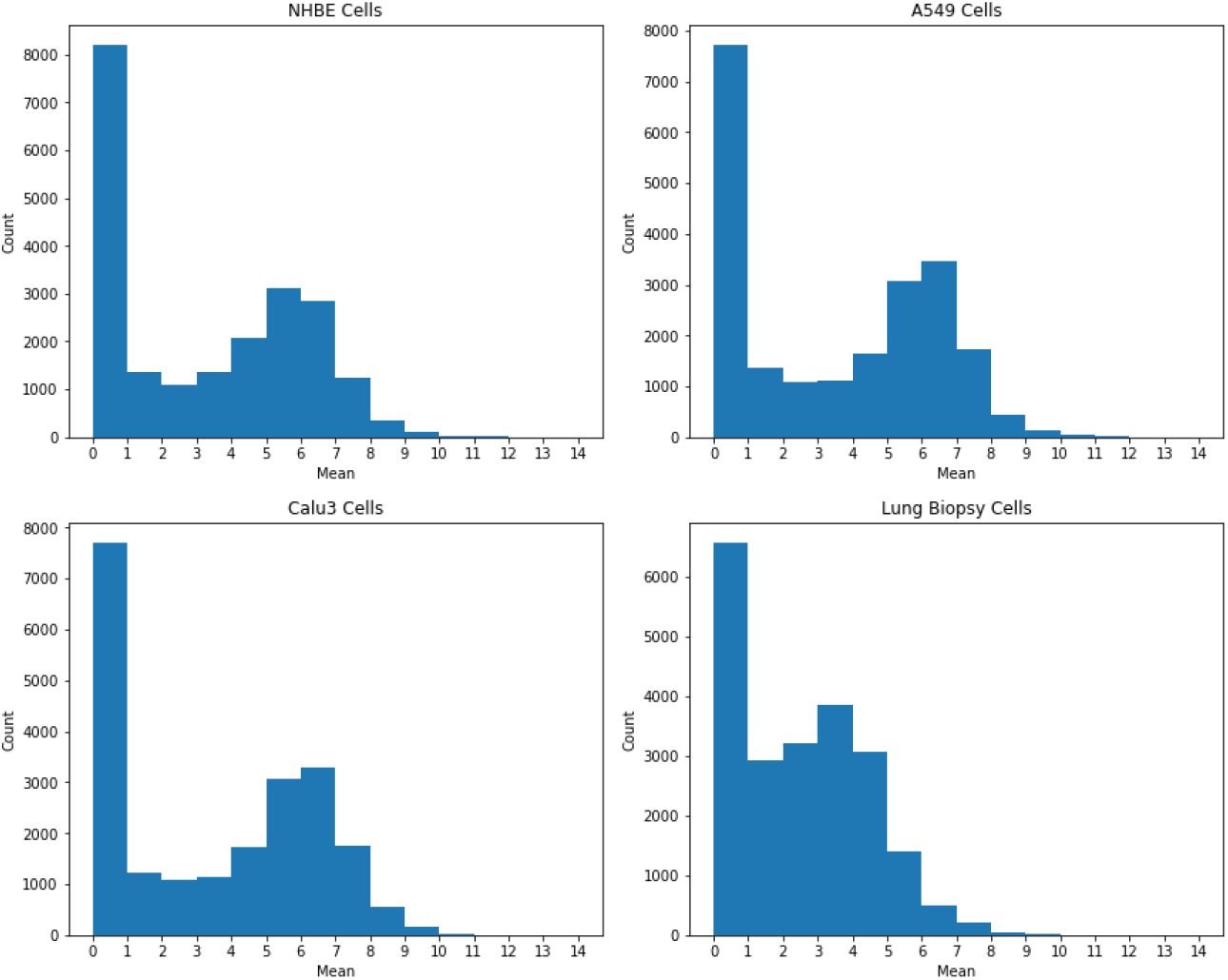
Distribution of gene expression (mean among samples) after applying a log2 transform (N = 21797 genes).

**Figure 3:**
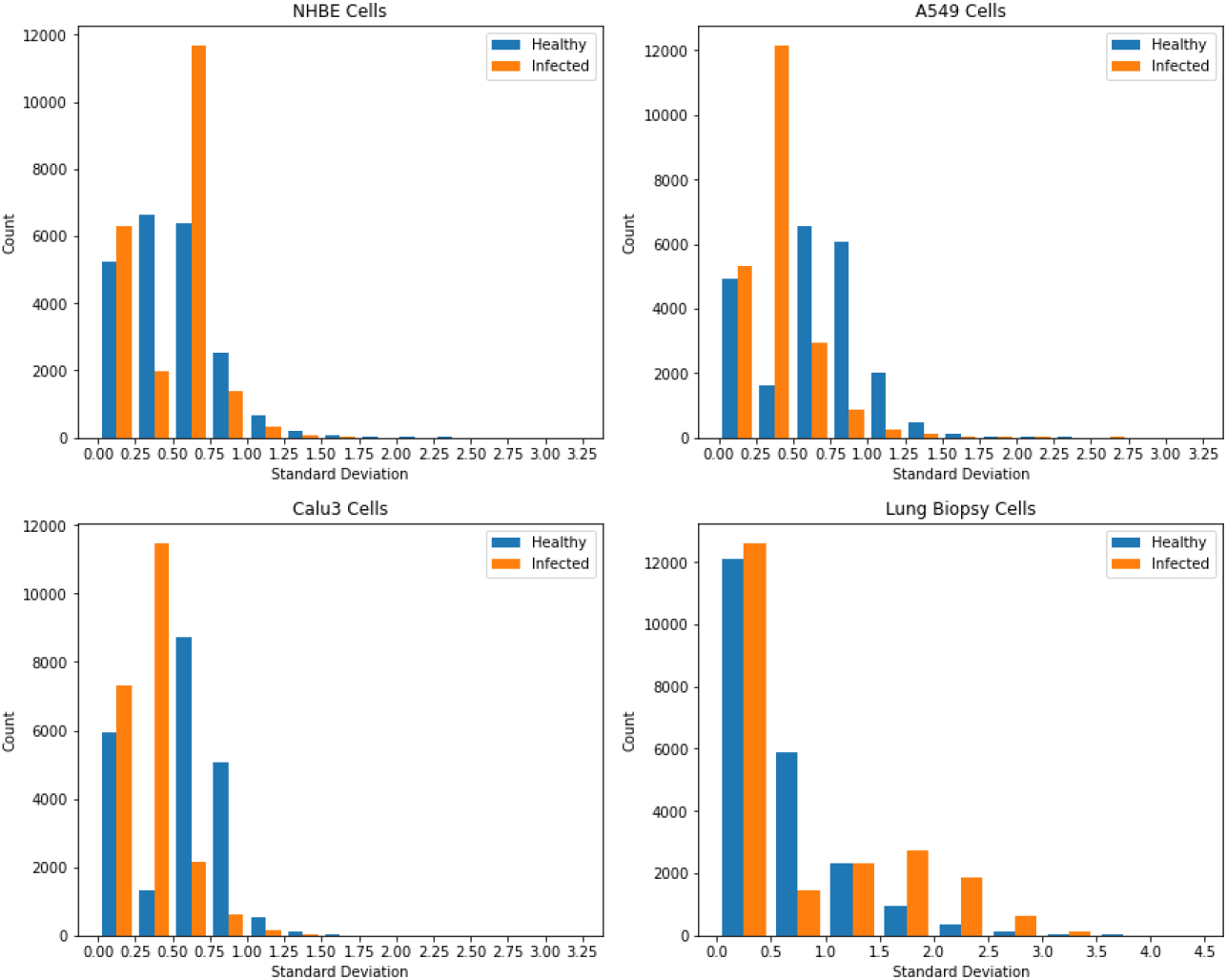
Standard deviation of gene expression within healthy and within infected cells.

In order to select an appropriate statistical test for the initial feature selection, a number of assumptions needs to be assessed. Firstly, we perform a median based Levene’s test [11], which is used, in the context of this work, to assess the equality of variances each pair of conditions (in particular for the pairs presented in Table 1). For these pairs, out of 19967 genes with non-null expression levels, 18990 had unequal variance for at least one pair of conditions, with *p <* 0.01. Additionally, a Shapiro-Wilk test [12] is used to assess whether these genes follow a normal distribution, applied in this case only to healthy and SARS-CoV-2 infected cells for each cell type (since these will be the main focus of our analysis and this test is only defined for at least 3 samples), with *p <* 0.05. It is important to note that overall 32.8%, 46.1% and 27.4% of genes for NHBE, A549 and Calu3 cells respectively are non-normal.

**Table 1:** Tested pairs of conditions.

| First Condition | Second Condition |
| --- | --- |
| NHBE Healthy | NHBE SARS-CoV-2 |
| NHBE Healthy | NHBE IAV |
| NHBE Healthy | NHBE IAVdNS1 |
| A549 Healthy | A549 SARS-CoV-2 |
| A549 Healthy | A549 IAV |
| A549 Healthy | A549 RSV |
| A549 Healthy | A549 HPIV3 |
| Calu3 Healthy | Calu3 SARS-CoV-2 |
| Biopsy Healthy | Biopsy SARS-CoV-2 |

The results of Levene’s test suggest that an assumption of equal variance cannot be made. As such, either an unequal variance (Welch) t-test or it’s non-parametric alternative, the Mann-Whitney U test [13], are more suitable for variable selection. With the results for non-normality still including a significant percentage of genes, Mann-Whitney U test is used.

## 4. Methods

The present work aims to find relevant biological processes involved in the infection of cells by SARS-CoV-2 using discriminative transcriptional modules produced from the application of state-of-the-art machine learning. To this end, we propose a methodology for the selection and discovery of correlated groups of differentially expressed genes (DEG) composed of five major steps. First, preprocessing techniques and preliminary gene selection are undertaken. Then, we proceed to pattern detection techniques, namely clustering, predictive modeling and biclustering. For each of these techniques, we apply functional enrichment to the obtained groups of genes in order to identify related biological functions. Finally, we analyse and interpret the identified functions, relating them to known characteristics of the disease as well as work by other authors. These steps are summarized in Figure 4.

**Figure 4:**
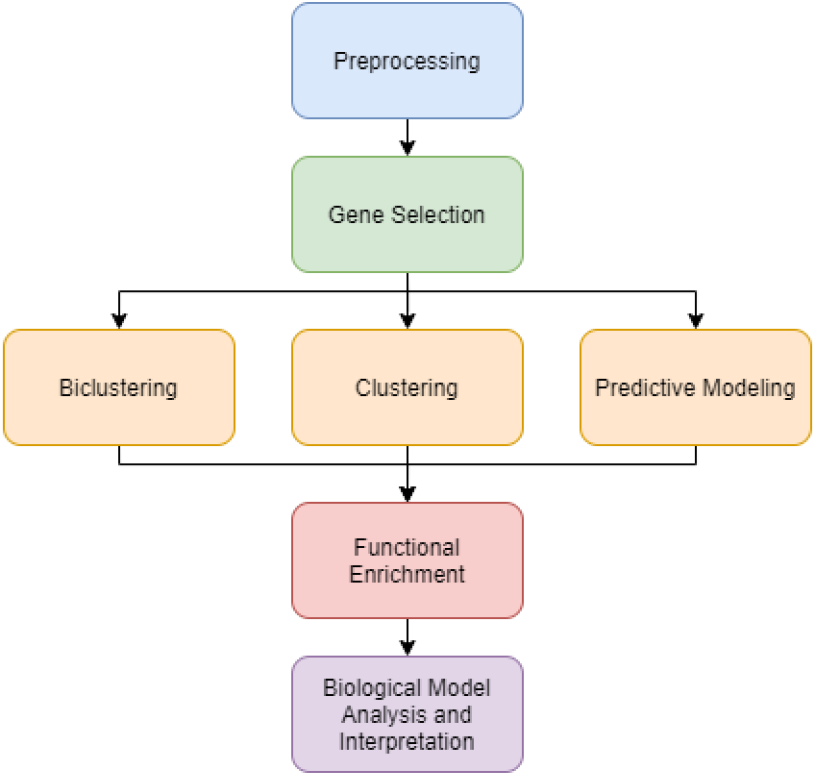
Schematic diagram of the steps composing the proposed computational pipeline.

**Figure 5:**
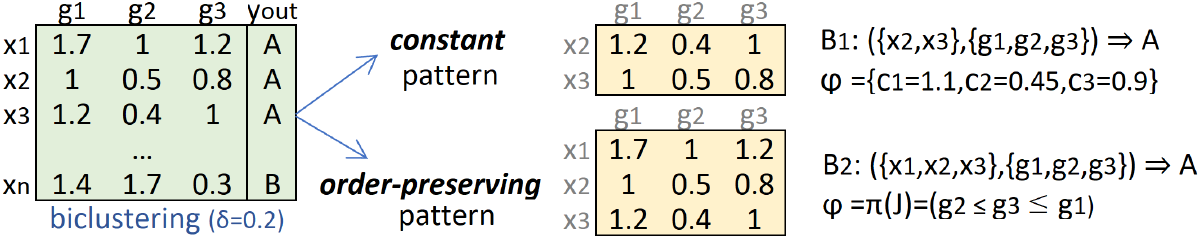
Biclustering with constant and order-preserving coherence assumptions. Constant bicluster has pattern (value expectations) *φ*_*B*_={*c*_1_=1.1, *c*_2_=0.45, *c*_3_=0.9} on *x*_2_ and *x*_3_ observations, while the order-preserving bicluster satisfies the *g*_1_ ≥ *g*_3_ ≥ *g*_2_ permutation on {*x*_1_, *x*_2_, *x*_3_} observations.

### 4.1. Preprocessing and Gene Selection

Given the highly skewed distribution of the data (with a vast majority of genes having low expression level), we first apply a log2 transform. Then, from the high-dimensional set of over 20.000 genes, we select a subset of DEG to be analysed. Due to the non-normal nature of data and unequal variance between control and test groups (as seen in section 3), Mann-Whitney U test is applied with a *p <* 0.05 and *p <* 0.01. By default a *p <* 0.01 is used, however for certain cell types *p <* 0.05 is suggested to guarantee a better coverage of gene candidates to the subsequent learning stage. The Mann Whitney U test tests for the null hypothesis that the two populations tested are equal. Therefore, this test can only be applied for pairs of conditions. We define the following settings, in which this method is applied:

- *Paired setting* : single pairs of conditions, e.g. healthy and SARS-CoV-2 infected NHBE cells or healthy and IAV infected A549 cells;
- *Multi-condition setting*: a set of pairs of conditions presented in Table 1. For each set of pairs, a Mann-Whitney U test is applied for each gene. Genes satisfying *p <* 0.01 or *p <* 0.05 are selected.

Additionally, ANOVA test [14] can be optionally applied to further ensure the discriminative power of the resulting differentially expressed genes. This is suggested if the subsequent learning step benefits from a space of reduced dimensionality by requiring to satisfy two distinct statistical criteria.

### 4.2. Pattern Detection

The usage of complete data with a simple statistical pre-selection of genes yields results which, depending on the chosen level of statistical significance, can surpass 1.000 genes. Applying functional enrichment to these results delivers none or very few enriched processes, which, when present, tend to be very generic cell functions. In this context, the pursue of putative transcriptional modules given by smaller sets of DEG is attempted to obtain more specific biological processes, as well as better statistical significance for each one found. To achieve this goal, three major approaches are applied: *clustering, predictive modeling* and *biclustering*.

#### 4.2.1. Clustering

The notion of cluster in our data can assume two distinct forms. First, a subset of correlated genes along a given set of samples. Second, a subset of correlated samples along a given set of genes. The latter is mainly interesting to understand which samples may be more closely related, though since it does not subdivide genes it cannot identify which genes may be better at distinguishing between different conditions. The former is thus suggested to identify sets of co-expressed genes. Agglomerative clustering is considered in this work with Euclidean affinity and Ward linkage. This is due to two main reasons: the easy visualization of the proximity between genes using a dendrogram, which can also help in the selection of the number of clusters; and the flexibility of the algorithm, which allows for multiple parameters to be adjusted according to the provided data. Despite the relevance of clustering for enrichment analysis [15, 16], it has considerable limitations. Namely, each gene set assesses expression similarly along all samples, which means, if multiple conditions are used simultaneously, this information will not be taken into account and will bias the detected patterns. However, by selecting different sets of conditions for each run of the algorithms, we can obtain relevant patterns for each specific condition and, though this doesn’t allow for a direct comparison between different conditions, it can provide sets of correlated genes which may have biological relevance.

#### 4.2.2. Machine learning models for classification

Classifiers generally use training data to produce predictive models, which are then used on test data to classify samples. In our work, since we seek to better understand potential signaling pathways and gene ontologies involved in the infection by SARS-CoV-2, we mainly focus on which genes are chosen to classify each of the samples, by inspecting the learned model. For this reason, we used associative classifiers given their easier explainability, namely decision trees [17], random forests [18] and XGBoost [19] in Python. While not directly interpretable, both random forests and XGBoost provide a metric of the relevance of each gene, which can be used to obtain the set of genes with the highest difference in expression level. In both cases, this metric corresponds to the impurity-based feature importances, which are calculated using the Gini criterion and then averaged across all trees within the model.

#### 4.2.3. Biclustering

Given a set of observations (conditions), *X*={*x*_1_, .., *x*_*n*_}, genes *G*={*g*_1_, .., *g*_*m*_}, a bicluster, *B*=(*I, J*), is a subspace defined by a subset genes, *J* ⊆ *G*, co-expressed on a subset of conditions, *I* ⊆ *X*. The *biclustering* task aims at identifying a set of biclusters, B, such that each bicluster, *B*_*k*_=(*I*_*k*_, *J*_*k*_), satisfies specific criteria of *homogeneity, dissimilarity* and *statistical significance*.

*Homogeneity* criteria are commonly guaranteed through the use of a merit function, such as the variance of the values in a bicluster [20]. Merit functions are typically applied to guide the formation of biclusters in greedy and exhaustive searches. In stochastic approaches, a set of parameters that describe the biclustering solution are learned by optimizing a merit (likelihood) function. The pursued homogeneity determines the coherence (co-expression patterning), quality (noise tolerance) and structure (number, size and positioning) of the subspaces in the biclustering solution [21]. A putative regulatory module is in this context given by a subspace of co-expressed genes, i.e. expression pattern on observations (Fig.5). A co-expressed subspace, *B*=(*I, J*), can be described by an *order-preserving coherence* when the ordering of a subset of genes according to their expression values, *π*_*J*_, is preserved for each sample in *I*. In alternative, co-expression can be defined by *constant coherence* where expression values in a bicluster, *a*_*ij*_ ∈ *B*, are described by *a*_*ij*_=*c*_*j*_+*η*_*ij*_, where *c*_*j*_ is the expected value of gene *g*_*j*_ and *η*_*ij*_ is the noise factor, generally a bounded deviation from expectations, *η*_*ij*_ ∈ [−*δ/*2, *δ/*2].

In addition to homogeneity criteria, *dissimilarity* criteria can be further placed to guarantee the discovery of non-redundant biclusters [22]. Finally, *statistical significance* criteria guarantee that the probability of a bicluster’s occurrence (against a null data model) deviates from expectations [23].

With biclustering algorithms, we can detect gene sets co-expressed on particular subsets of conditions, allowing for a more comprehensive modular view of regulatory responses to infection by SARS-CoV-2 and other viruses. In particular, when compared to the other proposed methods, biclustering allows for the detection of more specific patterns, such as a set of genes with higher or lower expression levels for a particular set of conditions, which are in turn easier to interpret and provide better results with functional enrichment.

To this end, we tested several algorithms to assess differences between the detected biclusters, namely the Cheng and Church [24], plaid [25] and xMotifs [26] algorithms. In recent years, a clearer understanding of the synergies between biclustering and pattern mining paved the rise of a new class of bi-clustering algorithms, generally referred to as *pattern-based biclustering* [21]. Pattern-based biclustering algorithms are inherently prepared to efficiently find exhaustive solutions of biclusters and offer the unprecedented possibility to affect their structure, coherency and quality [27]. This behavior explains why this class of biclustering algorithms are receiving an increasing attention in recent years [21]. In this context, we additionally assess the role of BicPAMS (Biclustering based on PAttern Mining Software), which consistently combines state-of-the-art contributions on pattern-based biclustering [22].

### 4.3. Functional Enrichment and Biological Analysis

To obtain potential biological processes associated with the gene groups found using the aforementioned methods, we used the EnrichR tool [28, 29]. To assess enrichment of terms in the target knowledge bases, we focus on three major criteria: p-value from Fisher’s exact test; the q-value, which adjusts the p-value to control the False Discovery Rate; and the z-score, which takes into account that Fisher’s exact method to calculate the *p*-value produces lower values for longer lists even if they are random. Furthermore, we also consider the combines the z-score and the p-value as follows: *c* = ln(*p*) × *z*. We prioritize both the adjusted *p*-value and the combined score to compare the results of the enrichment analysis.

Additionally, EnrichR web tool provides access to multiple knowledge bases (https://maayanlab.cloud/Enrichr/libraries). For our analysis, we prioritize Gene Ontology (GO) Biological Process knowledge base [30, 31] (ver. 2021) as it comprehensively characterizes a large amount of genes (14937) against 6036 terms, including recently augmented biological processes on viral infection and immune responses. Additionally, we use the Kyoto Encyclopedia of Genes and Genomes (KEGG) [32] to analyse enriched pathways and diseases. The identified biological processes are then analysed and compared to known characteristics of the disease and other previous studies, in order to identify potential new insights into the effects of the virus and verify existing ones.

### 4.4. Code Availability

The code used to obtain the results can be obtained in the following GitHub repository: https://github.com/PRodrigues98/Analysis-of-regulatory-response-to-SARS-CoV-2-infection. Dependencies: Python version 3.8, NumPy, pandas, scikit-learn and matplotlib libraries.

## 5. Results and Discussion

To address the challenges of classic functional enrichment analyzes, we considered three approaches for identifying DEGs associated with modular regulatory views as highlighted in section 4: clustering, predictive modeling and biclustering. In the present section, we present the key findings resulting from the application of each of these methods to the dataset, as well as an analysis of the identified biological processes within the context of viral infection. In particular, we will begin with clustering, then classification, and finally biclustering.

To assess the effectiveness of the methods, we begin by presenting, in Table 2, the result of performing functional enrichment on the baseline set of genes obtained directly through preprocessing (using the Multi-Condition Setting, *p <* 0.01 in accordance with section 4). As we can see in Table 2, there is a considerable number of processes with low p-value. However, the c-score is significantly lower when compared to the same genes after clustering (see Table 2). This is likely due to the higher number of genes being analysed together when compared to the proposed methods, since clustering and biclustering identify smaller subgroups of genes with correlated expression and predictive models select a smaller number of genes. Additionally, terms such as *negative regulation of bone remodeling* (GO:0046851) and *negative regulation of bone resorption* (GO:0045779), which seem to be more generic and less related to the viral infection appear in this analysis, but do not seem to reoccur within the terms found for clustering, classification or biclustering.

**Table 2:** Top 8 GO biological processes ordered by combined score after single preprocessing (Multi-Condition Setting ^4^, *p <* 0.01).

| GO Biological Process | p-value | c-score |
| --- | --- | --- |
| cellular response to type I interferon (GO:0071357) | 2.35E-10 | 324.03 |
| type I interferon signaling pathway (GO:0060337) | 2.35E-10 | 324.03 |
| cytokine-mediated signaling pathway (GO:0019221) | 3.80E-26 | 319.38 |
| protein mono-ADP-ribosylation (GO:0140289) | 3.22E-04 | 319.17 |
| receptor signaling pathway via STAT (GO:0097696) | 2.73E-06 | 299.82 |
| receptor signaling pathway via JAK-STAT (GO:0007259) | 2.50E-06 | 250.15 |
| exogenous peptide antigen, TAP-independent (GO:0002480) | 7.04E-03 | 219.44 |
| negative regulation of bone remodeling (GO:0046851) | 2.62E-03 | 212.68 |

### 5.1. Clustering analysis

Considering the application of clustering in the Multi-Condition Setting (*p <* 0.01) in Table 3, a high percentage of the top identified processes are related to response to viral infection, as well as to immune responses. The annotation *cytoplasmic pattern recognition receptor (PRR) signaling pathway in response to virus*, GO:0039528 (directly related to the annotations GO:0140546 and GO:0051607, also within the top 25 enriched processes) corresponds to a set of molecular signals associated with the detection (by binding of viral RNA molecules to certain cytoplasmic receptors) of a virus. In particular, the detection seems to be performed by the RIG-I PRR, responsible for the detection of RNA synthesized during the process of viral replication, since there are 3 child processes (GO:0039529 with *p* = 2.91× 10^*−*3^ and *c* = 905.29; GO:0039535 with *p* = 7.67 × 10^*−*4^ and *c* = 526.08; GO:0039526 with *p* = 5.26 × 10^*−*3^ and *c* = 513.51) associated with this receptor which are still statistically relevant. This receptor, along with others, has been identified as part of the inflammatory response to SARS-CoV-2 as well as other coronaviruses [33]. Additionally, the signaling cascade resulting from the detection of viral proteins is associated with the production of Type I interferons and pro-inflammatory cytokines [34], which can also be observed within the top enriched processes (for instance, terms GO:0060337, GO:0071357 and GO:0060333).

**Table 3:** Top 8 GO biological processes ordered by combined score (Multi-Condition Setting, Clustering, *p <* 0.01).

| GO Biological Process | p-value | c-score |
| --- | --- | --- |
| type I interferon signaling pathway (GO:0060337) | 4.40E-27 | 9111.11 |
| cellular response to type I interferon (GO:0071357) | 4.40E-27 | 9111.11 |
| negative regulation of viral genome replication (GO:0045071) | 1.10E-16 | 3820.31 |
| defense response to symbiont (GO:0140546) | 7.74E-22 | 3260.22 |
| cytoplasmic PRR signaling pathway <sup>5</sup>(GO:0039528) | 1.74E-06 | 3253.53 |
| negative regulation of viral process (GO:0048525) | 5.16E-17 | 3163.10 |
| defense response to virus (GO:0051607) | 2.34E-21 | 2930.10 |
| endogenous peptide antigen, TAP-independent (GO:0002486) | 4.42E-05 | 2797.75 |

In particular, the term *type I interferon signaling pathway* (GO:0060337), which has several related terms also present within the top 25 processes (for instance, *type I interferon signaling pathway*, GO:0060337 and *cytokine-mediated signaling pathway*, GO:0019221, both direct ancestors) are related to type I interferons. The association between these and the process of viral infection is further bolstered by the presence of terms *response to interferon-beta* (GO:0035456) and *response to interferon-alpha* (GO:0035455), which are both type I interferons.

It is also interesting to note the presence of the term *negative regulation of type I interferon-mediated signaling pathway* (GO:0060339) as well as *negative regulation of chemokine production* (GO:0032682). Chemokines are involved in inflammation and the control of viral infections, and they and their receptors are sometimes mimicked by viruses in order to evade host antiviral immune responses [35]. The presence of these is noteworthy mainly due to directly opposing the other processes related to the activation of an immune response.

Additionally, there are multiple processes directly related to cellular response to viruses, namely *defense response to symbiont* (GO:0140546), *defense response to virus* (GO:0051607), *negative regulation of viral genome replication* (GO:0045071, also associated with GO:0045069), *antiviral innate immune response* (GO:0140374), *negative regulation of viral process* (GO:0048525) and *cellular response to virus* (GO:0098586). These indicate that NHBE cells were able to identify that they had been infected by a virus and induce an immune response.

For A549 cells, the genes composing all detected processes have higher expression levels for infected cells than for control. Similarly to NHBE cells, there seems to be a prevalence of type I interferon and cytokine related terms. Multiple processes, such as *cellular response to type I interferon* (GO:0071357), *type I interferon signaling pathway* (GO:0060337), *response to interferon-beta* (GO:0035456) are repeated, with most of the common processes having to do with interferon and general cytokine response as well as responses to viral infection.

The terms *STAT cascade* (GO:0097696), *positive regulation of JAK-STAT cascade* (GO:0046427) and *JAK-STAT cascade* (GO:0007259), are not present for NHBE cells. These are all related to the JAK-STAT signaling pathway, which is associated with a wide variety of cytokines. Not triggering signaling or not regulating it properly, can lead to inflammatory disease [36], among other issues.

Interestingly, similarly to the NHBE cells the process *negative regulation of type I interferon production* (GO:0032480) seems to suggest a potential attempt to reduce immune response. However, the opposite term, *positive regulation of type I interferon production* (GO:0032481) is also within the top 25 (though with higher p-value and lower c-score). This may be due to both pathways being active simultaneously, although it may also reveal overlap in the genes that produce each process (2 out of 5 genes in common between the two processes).

In Table 4 we present the pathways identified when using the KEGG Pathway database (ver. 2021). The results, similarly to the GO database, include multiple virus related pathways. These are all composed by genes with higher expression values for infected cells than control. Within the top identified terms, there is a prevalence of virus related pathways. *Coronavirus disease* (map05171), *the sixth term, is directly associated with SARS-CoV-2, which provides more confidence that the terms identified thus far are indeed related to the viral infection*.

**Table 4:** Top 8 KEGG pathways ordered by combined score (A549 cells, healthy vs SARS-CoV-2).

| KEGG Pathway | p-value | c-score | Cluster |
| --- | --- | --- | --- |
| Measles (map05162) | 1.02E-08 | 295.14 | 2 |
| Influenza A (map05164) | 1.02E-08 | 252.18 | 2 |
| Herpes simplex virus 1 infection (map05168) | 7.16E-10 | 177.20 | 2 |
| Epstein-Barr virus infection (map05169) | 4.82E-07 | 156.14 | 2 |
| TNF signaling pathway (map04668) | 1.11E-07 | 153.86 | 0 |
| Coronavirus disease (map05171) | 3.78E-07 | 150.32 | 2 |
| RIG-I-like receptor signaling pathway (map04622) | 3.68E-04 | 120.34 | 2 |
| NOD-like receptor signaling pathway (map04621) | 9.37E-06 | 119.63 | 2 |

Additionally, the term RIG-I*-like receptor signaling pathway* (map04622), which is related to the previously mentioned RIG-I receptor, helps solidify the idea of it being involved in the anti-viral immune response.

The KEGG pathways *Antigen processing and presentation* (map04612), *JAK-STAT signaling pathway* (map04630), the principal signaling mechanism for a variety of cytokines, *IL-17 signaling pathway* (map04657), a subset of cytokines with various roles related to inflammatory responses and defence against external pathogens, and *NF-kappa B signaling pathway* (map04064), a signaling pathway which is activated by the aforementioned cytokines and is related to immune responses, all support the processes identified previously in the role played by inflammatory cytokines and related signaling pathways in the infection by SARS-CoV-2.

### 5.2. Predictive modeling analysis

For predictive models, we considered the top relevant genes selected by the Random Forest and xGBoost algorithms. With XGBoost, 94 genes are identified. With the Random Forest, 356 genes are selected. These algorithms have 69 genes in common. There are multiple terms present in both models, mostly related to immune system activity. However, there are several processes uniquely identified by each of the algorithms. Processes identified only by XGBoost are particularly interesting, since most genes selected by XGBoost are also selected by the Random Forest and the extra genes selected by the Random Forest may mask relevant information.

*ISG15-protein conjugation* (GO:0032020), a term identified only within XGBoost selected genes, is related to the cellular protein modification process of ISG15. This protein has an important role in host antiviral response, with several different actions depending on the infecting virus. Most significantly among these actions is the inhibition of viral replication in addition to the modulation of the damage and repair as well as the immune responses [37].

Also within the terms identified only by XGBoost, there are multiple related to chemotaxis, the movement of a cell or organism towards a higher or lower concentration of a given substance, and migration of various types of immune cells. In particular, macrophages [38, 39] (GO:0048246 and GO:1905517), natural killer cells [40] (GO:2000501)), eosinophils [41] (GO:0072677 and GO:0048245), neutrophils [42] (GO:0030593 and GO:1990266), which are all types of white blood cells involved with the innate immune response to viral infection.

Additionally, there are multiple terms in both cases associated with cytokine production and related signaling pathways, as well as response to different types of interferons. In addition to these, terms such as *regulation of fever generation* (GO:0031620), *negative regulation of viral process* (GO:0048525), *inflammatory response* (GO:0006954) and *negative regulation of viral genome replication* (GO:0045071) are also associated with immune response. Together with the previously mentioned signaling of white blood cells, these results show the significant, both innate and adaptive, immune responses by cells infected by this virus.

Among the top processes in both tables is *chronic inflammatory response* (GO:0002544). Similarly to what was mentioned for the combined data, there are multiple terms related to the recruitment of certain types of white blood cells. In particular, *positive regulation of monocyte chemotactic protein-1 production* (GO:0071639), the top term for the Random Forest, is associated to a protein which plays a key role in the migration of monocytes [43].

It is also important to note that multiple terms associated with the apoptotic process are present, namely *positive regulation of intrinsic apoptotic signaling pathway* (GO:2001244), *regulation of intrinsic apoptotic signaling pathway* (GO:2001242) and *positive regulation of apoptotic signaling pathway* (GO:2001235). This process, responsible for causing the death of a cell when a certain internal or external stimulus is received, may indicate that the cell was able to detect that it was infected by SARS-CoV-2. This hypothesis is further supported by the presence of the term *pattern recognition receptor signaling pathway* (GO:0002221). These receptors, as previously explained for the related term present in Table 3, have been associated with the inflammatory response to SARS-CoV-2 [33].

There are several terms related to the response to virus by the host. In particular, *positive regulation of defense response to virus by host* (GO:0002230), *regulation of defense response to virus by host* (GO:0050691), *defense response to symbiont* (GO:0140546) and *defense response to virus* (GO:0051607), although these are only present within the Random Forest selected genes. It is also worth noting once again the abundance of interferon related processes, as well as some cytokine related terms. Among these, *negative regulation of cytokine production* (GO:0001818) and *positive regulation of cytokine production* (GO:0001819), which are contradicting, may indicate an attempt to modulate the immune response by the cell or potentially a mechanism of the virus to defend itself from the immune response.

The term *RIG-I signaling pathway* (GO:0039529) which is associated with the Pattern Recognition Receptor RIG-I, and the term *cytoplasmic pattern recognition receptor signaling pathway in response to virus* (GO:0039528) were also identified in Table 3 as well as for NHBE cells using the Random Forest algorithm. These receptors play crucial roles in the detection of viruses by cells and the resulting signaling cascade, which in turn leads to the production of Type I interferons and pro-inflammatory cytokines [34].

### 5.3. Biclustering analysis

In order to allow for the detection of more complex patterns, we now present the results of applying several biclustering algorithms to our data. In particular, these algorithms, unlike clustering, can identify patterns which span only certain conditions. This means that by analyzing the resulting bi-clusters and functionally enriching them, we can obtain processes associated with any particular subset of conditions.

We begin in Table 5 by presenting several metrics for each algorithm and preprocessing option used. BicPAMS and Cheng and Church present the highest average number of biclusters, with the Plaid and xMotifs algorithms significantly less for most preprocessing conditions. It is also important to note that BicPAMS presents delineatedly higher enrichments, and selects a larger amount of genes for a much smaller amount of conditions. Comparatively, this behavior is particularly relevant since having too many conditions can lead to the identification of more generic genes and having too few genes can lead to the identification of less significant processes.

**Table 5:** Metrics for comparing the performance of the tested biclustering algorithms with <u>di</u>fferent preprocessing techniques. Note: |ℬ| corresponds to the number of biclusters; 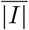 is the average number of genes per bicluster; 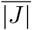 the standard deviation of genes per bicluster; |*J*| the average number of conditions per bicluster; *σ*_|*J*|_ the standard deviation of the number of conditions per bicluster; and finally 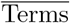 the average number of enriched terms per bicluster.

| Algorithm | Preprocessing | $\vert \mathcal{B}\vert $ | $\overline{\vert I\vert }$ | $\sigma_{\vert I\vert }$ | $\overline{\vert J\vert }$ | $\sigma_{\vert J\vert }$ | $\overline{\text{Terms}}$ |
| --- | --- | --- | --- | --- | --- | --- | --- |
| BicPAMS | $p < 0.01$ | 80 | 208.03 | 18.54 | 3.16 | 0.53 | 28.91 |
| | $p < 0.05$ | 79 | 3526.66 | 301.50 | 3.24 | 0.64 | 341.70 |
|  | ANOVA (top 200) | 7 | 188.29 | 5.95 | 10.00 | 9.70 | 10.57 |
|  | ANOVA (top 1000) | 20 | 676.05 | 29.13 | 5.00 | 4.22 | 55.75 |
|  | ANOVA (top 5000) | 57 | 2106.18 | 128.36 | 3.61 | 1.25 | 131.32 |
| Cheng and Church | $p < 0.01$ | 50 | 15.60 | 12.59 | 12.92 | 5.90 | 3.46 |
| | $p < 0.05$ | 100 | 55.90 | 16.23 | 34.79 | 9.96 | 1.68 |
|  | ANOVA (top 200) | 8 | 25.00 | 23.49 | 21.38 | 12.56 | 6.50 |
|  | ANOVA (top 1000) | 56 | 17.86 | 15.27 | 17.89 | 10.67 | 4.41 |
|  | ANOVA (top 5000) | 100 | 34.54 | 24.10 | 22.76 | 11.25 | 2.47 |
| Plaid | $p < 0.01$ | 10 | 64.70 | 55.53 | 14.20 | 5.60 | 29.90 |
| | $p < 0.05$ | 10 | 776.40 | 922.43 | 11.60 | 6.89 | 24.10 |
|  | ANOVA (top 200) | 8 | 44.00 | 30.76 | 12.88 | 8.43 | 9.88 |
|  | ANOVA (top 1000) | 10 | 159.50 | 100.72 | 12.20 | 7.29 | 18.70 |
|  | ANOVA (top 5000) | 10 | 739.20 | 530.04 | 13.10 | 7.48 | 43.40 |
| xMotifs | $p < 0.01$ | 10 | 31.90 | 17.17 | 8.20 | 2.86 | 1.70 |
| | $p < 0.05$ | 10 | 654.50 | 365.82 | 6.00 | 0.00 | 6.30 |
|  | ANOVA (top 200) | 6 | 30.33 | 34.30 | 24.50 | 9.73 | 10.67 |
|  | ANOVA (top 1000) | 10 | 71.90 | 103.54 | 11.10 | 4.28 | 7.60 |
|  | ANOVA (top 5000) | 10 | 326.00 | 538.95 | 6.20 | 0.60 | 5.30 |

Using these methods, we obtain a set of biclusters, each consisting of a subset of genes and a subset of conditions. By performing functional enrichment on these genes, a set of biological processes associated with those genes is then produced. In order to analyze these results and obtain a more generic view of how often certain processes occur for each condition, a count is performed for each process identified. This allows for the identification of the most commonly occurring processes, and thus provides a better view of which processes are most closely related with a certain condition, while also potentially reducing the amount of more generic biological processes. In addition to this, it provides a direct element of comparison between different cell types for the same condition, or between the same cell type and different viruses. In addition to the number of occurrences of each process, the best c-score and p-value are also provided, in order to compare the statistical relevance of different processes.

We now proceed to a comparative analysis of the biological processes associated with SARS-CoV-2 for all cell types, using biclustering. In order to provide an ordering for the processes taking into account all cell types, each enriched term is first ranked by the number of occurrences it has related to a given condition. Then a fused rank is computed by multiplying the resulting ranks. The multiplication allows for a higher penalization of terms which contain a single very low rank but high ranks for other cell types.

There are several identified processes which have been previously described with clustering and predictive models. In particular, there are multiple terms related to cytokine activity, for instance *cytokine-mediated signaling pathway* (GO:0019221), which possesses a high number of occurrences for A549 (1.00), NHBE (0.75) and Calu3 (1.00) cells and a lower count for Biopsy cells (0.60). It is interesting to note a seeming tendency for the normalized number of occurrences for Biopsy cells to be lower for most processes, with more generic DNA related processes, such as *DNA metabolic process* (GO:0006259), *DNA repair* (GO:0006281) and *cellular response to DNA damage stimulus* (GO:0006974), possessing higher values. This may be due to biopsy results possibly containing multiple cell types as well as due to the very low number of samples of this type of cell (2 healthy and 2 infected).

Other cytokine associated processes include *cellular response to cytokine stimulus* (GO:0071345), *chemokine-mediated signaling pathway* (GO:0070098) followed also by *cellular response to chemokine* (GO:1990869). Chemokines in particular play an important role in multiple processes related with host immune response against viral infection, namely the attraction of leukocytes to the infected tissue. The presence of the terms *neutrophil mediated immunity* (GO:0002446), *neutrophil activation involved in immune response* (GO:0002283) and *neutrophil degranulation* (GO:0043312), further supports this hypothesis. Neutrophils are leukocytes which are the first responders to sites of infection, and have also been identified as the main infiltrating cell population in IAV infection [42]. Despite containing somewhat lower counts than other processes, this set of enriched terms still possess p-values and c-scores well within the range of statistical significance.

Another previously identified set of processes which is also present is interferon related terms. Interferons are a potent type of cytokines which are associated with antiviral response, with most viruses having developed adaptations to at least partially avoid this mechanism [44]. In particular, *cellular response to interferon-gamma* (GO:0071346) and *interferon-gamma-mediated signaling pathway* (GO:0060333).

We now proceed to a comparative analysis of the processes associated with different viruses. There are many processes in common with the SARS-CoV-2 analysis, which is to be expected, since most identified processes are related to immune response.

*Cellular response to interferon-gamma* (GO:0071346) has somewhat fewer occurrences when compared to the other viruses (0.66 vs 0.85 for RSV, 0.92 for HPIV3 and 0.86 for IAV). *Cytokine-mediated signaling pathway* (GO:0019221) has a somewhat higher number of occurrences for SARS-CoV-2 and HPIV than others (1.00 and 1.00 vs 0.74 for RSV and 0.92 for IAV). *Inflammatory response* (GO:0006954) is somewhat muted for SARS-CoV-2 when compared to the other viruses, for both A549 (0.45 vs 0.97 for RSV, 0.92 for HPIV3, 1.00 for IAV) and NHBE cells (0.75 vs 0.91 for IAV, 1.00 for IAVdNS1). These differences are consistent with those found by Blanco-Melo D. et al. [1], who found SARS-CoV-2 to induce a limited interferon response when compared with the other viruses but a strong production of cytokines and resulting processes. Overall, there seems to be a tendency for the other viruses to have comparatively higher counts, especially IAV.

In Table 6, we can see a compilation of the number of GO Biological Processes detected for each of the applied methods. As we can see, biclustering provided, by a considerable margin, a highest amount of biological processes, followed by clustering. The predictive models generally provided worse results, with Random Forests providing somewhat better results amongst predictors for the Multi-Condition Setting as well as for NHBE cells. Overall, these results provide initial empirical evidence in favor of pattern-based algorithms to promote the coverage and statistical significance of functional enrichment analysis, offering a way of unraveling less-trivial yet relevant regulatory behavior in knowledge bases.

**Table 6:**
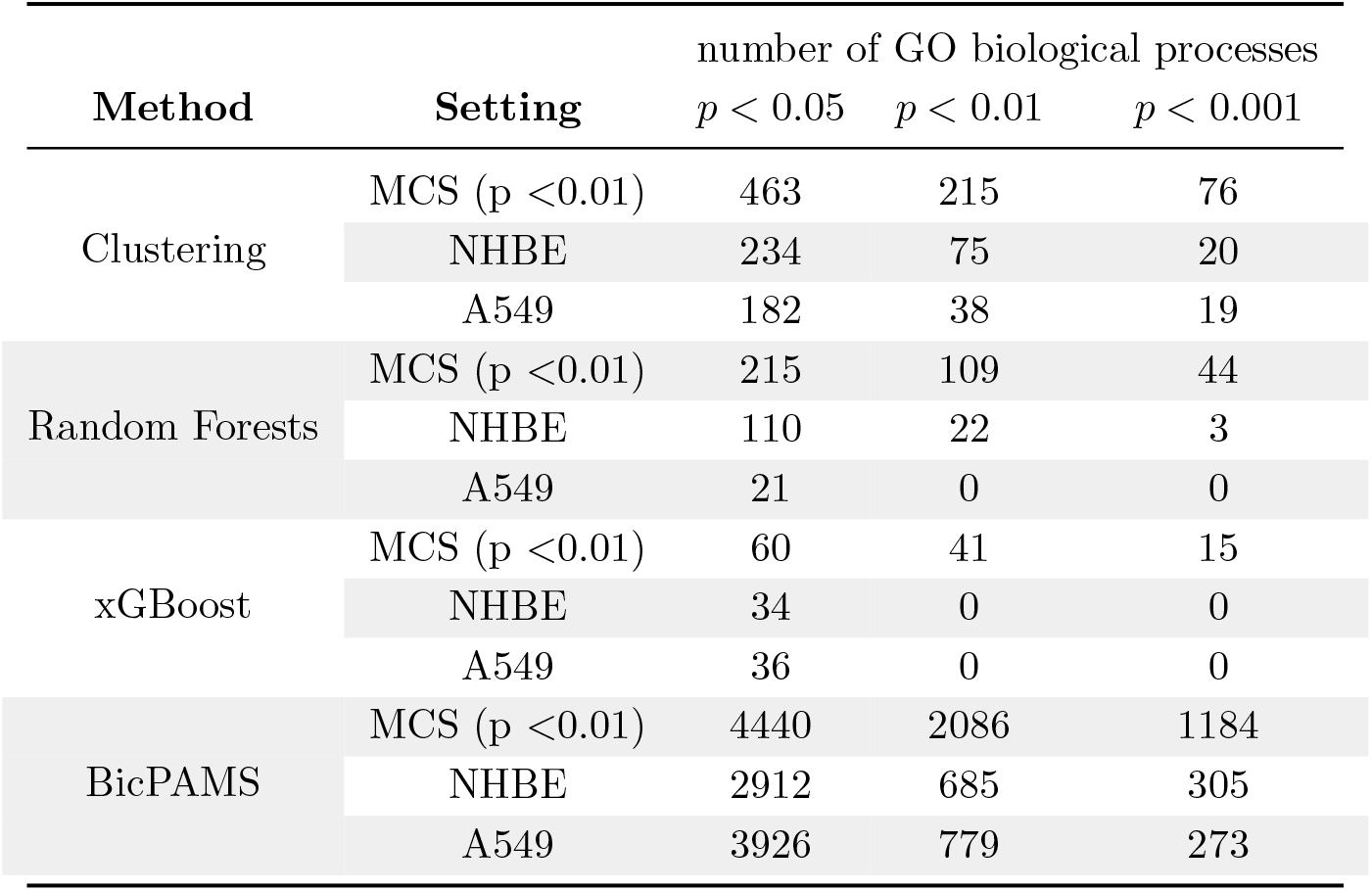
Number of processes found, for different *p* values, for each of the methods applied (Multi-Condition Setting).

## 6. Conclusion

This work assesses the impact of different regulatory modules views for gene set enrichment analysis from gene expression responses to SARS-CoV-2 infection, while also presenting an analysis of non-trivial biological processes associated with virus infection. A particular focus is placed on the relevance of pattern-centric views given the observed role of subspace clustering stances to improve the coverage and quality of enriched terms from knowledge bases.

A novel methodology is proposed, combining different approaches, which when consolidated provide a more robust view of the putative processes associated with virus infection. In particular, the complete gene set is initially filtered using a Mann-Whitney U Test, which allows for the selection of genes with statistically relevant differences in expression between healthy and infected cells. Other authors perform feature enrichment directly on the set of genes obtained using simplistic statistical tests. However, this stance results in a smaller amount of biological processes detected, as well as a decrease in their quality (measured using Fisher’s Exact Test and the combined c-score). So a three-fold, pattern-centric approach – composed by clustering, associative predictive modeling and biclustering algorithms – is suggested to identify DEGs with correlated expression.

Under this methodology, we were able to validate and identify potentially novel biological processes associated with SARS-CoV-2 infection. Among the various enriched terms, the high cytokine induction, Type I interferon related terms, as well as signaling pathways related to these were reoccurring in all analysis performed. In particular, SARS-CoV-2 was found to induce a limited interferon response when compared with the other viruses but a strong production of cytokines and associated processes (namely interferon induction and response to these stimuli). These findings were consistent as in the previous case study [1]. Additionally, we found in multiple analysis the involvement of Pattern Recognition Receptors (with particular emphasis on RIG-I) in the process of infection. This was not identified in previous studies, however it is consistent with other literature on coronaviruses, and further supports the hypothesis that a pattern-centric view of the gene enrichment process can result in novel information.

As directions for future work, we aim at: i) extending the conducted experimental analysis towards other transcriptomic data sources, in particular SARS-CoV-2 related sources, to cross-validate, expand and improve the robustness of the provided findings; ii) assessing the validity of the methodological contributions over proteomic and metabolomic data; iii) addressing the issue of sample interdependence by testing the underlying relationships and designing pattern-centric approaches tailored to the presence of replicates; and iv) explicitly combining available background knowledge [45] to guide the enrichment analysis towards novel findings.

Finally, we believe that this proposed computational pipeline will prove to be a useful workflow for generating hypotheses in other biological datasets.

## Funding

This work is supported by Fundação para a Ciência e Tecnologia (FCT) under IPOscore (DSAIPA/DS/0042/2018), LAETA (UIDB/50022/2020), and ILU (DSAIPA/DS/0111/2018). This work was further supported by LAQV, financed by national funds from FCT/MCTES (UIDB/50006/2020 and UIDP/50006/2020) and INESC-ID plurianual (UIDB/50021/2020). The FCT distinction (CEECIND/01399/2017) to RSC under the CEEC Individual program is also acknowledged.

## Footnotes

1 https://www.who.int/emergencies/diseases/novel-coronavirus-2019, accessed on Dec. 5th, 2021

2 Available at https://www.ncbi.nlm.nih.gov/geo/query/acc.cgi?acc=GSE147507

4 Previously defined in subsection 4.1, Table 1

5 some names have been shortened in favor of succinctness, with full definitions available in the accompanying hyperlink

## Notes

### Competing Interest Statement

The authors have declared no competing interest.

